# Development of CAR-T Cell Therapy for B-ALL Using a Point-of-Care Approach

**DOI:** 10.1101/770297

**Authors:** Luiza de Macedo Abdo, Luciana Rodrigues Carvalho Barros, Mariana Saldanha Viegas, Luisa Vieira Codeço Marques, Priscila de Sousa Ferreira, Leonardo Chicaybam, Martín Hernán Bonamino

**Affiliations:** Immunology and Tumor Biology Program – Research Coordination – Brazilian National Cancer Institute (INCA), Rio de Janeiro, Brazil; Vice-Presidency of Research and Biological Collections (VPPCB) – Oswaldo Cruz Foundation (FIOCRUZ), Rio de Janeiro, Brazil

## Abstract

Recently approved by the FDA and European Medicines Agency, CAR-T cell therapy is a new treatment option for B-cell malignancies. Currently, CAR-T cells are manufactured in centralized facilities and face bottlenecks like complex scaling up, high costs and logistic operations. These difficulties are mainly related to the use of viral vectors and the requirement to expand CAR-T cells to reach the therapeutic dose. In this paper, by using Sleeping Beauty-mediated genetic modification delivered by electroporation, we show that CAR-T cells can be generated and used without the need for *ex vivo* activation and expansion, consistent with a point-of-care (POC) approach. Our results show that minimally manipulated CAR-T cells are effective *in vivo* against RS4;11 leukemia cells engrafted in NSG mice even when inoculated after only 4 hours of gene transfer. In an effort to better characterize the infused CAR-T cells, we show that 19BBz T lymphocytes infused after 24h of electroporation (where CAR expression is already detectable) can improve the overall survival and reduce tumor burden in organs of mice engrafted with RS4;11 or Nalm-6 B cell leukemia. A side-by-side comparison of POC approach with a conventional 8-day expansion protocol using Transact beads demonstrated that both approaches have equivalent antitumor activity *in vivo*. Our data suggests that POC approach is a viable alternative for the generation and use of CAR-T cells, overcoming the limitations of current manufacturing protocols. Its use has the potential to expand CAR immunotherapy to a higher number of patients, especially in the context of low-income countries.

## Introduction

In the last decade, CAR-T cell immunotherapy has shown considerable results in the treatment of different types of cancer, especially of patients with B cell malignancies, with ~80% of response rate in patients with B cell acute lymphoblastic leukemia (1,2) and 52-82% response rate in patients with B cell lymphoma(3,4). This therapy consists of redirecting T lymphocytes to antigens expressed by tumor cells by expressing a chimeric receptor harboring an extracellular antigen recognition domain derived from an antibody, a transmembrane region and an intracellular domain for signaling and activation of the T lymphocyte (5). Based on these results, anti-CD19 CAR-T cells were recently approved by the FDA and European Commission for use in USA and Europe, consisting in a new treatment option for patients with B cell malignancies. Similar therapies targeting antigens like CD22(6,7), BCMA(8–10) and EGFRvIII(11) are currently being evaluated in clinical trials, which will further extend the clinical application of CAR-T cell therapy to other types of cancer and potentially improve the response rates in the diseases already approached by CAR-T cell therapies.

Currently, the prevailing CAR-T cell manufacturing process involves the use of retro or lentiviral vectors for delivering the transgene to purified T cells followed by an *in vitro* expansion protocol aimed at generating enough T lymphocytes to reach the target dose, ranging in general from 2-5×10^6^/kg(12). This process, despite providing satisfactory performance in generating the currently approved therapies, will hardly meet the expected increase in demand for CAR-T cell therapies in the near future, both in terms of cost and time of production. Retroviral and lentiviral vectors are costly and cumbersome to produce in large batches, and their use requires that specific quality control assays regarding the presence of replication-competent retrovirus (RCR) are performed in the final product(13). Moreover, use of retroviral vectors requires pre-activation of T cells, which generally adds at least 2 days to the manufacturing process. In combination with the current methods of T cell expansion, like Wave bioreactors, or G-REX flask, total production time ranges from 12 to 16 days(14).

We and others have shown that the integrative, non-viral Sleeping Beauty (SB) transposon system is a suitable alternative to viral vectors in the process of CAR-T cell production(15–18). CAR-T cells generated by electroporation of mononuclear cells with SB plasmids (one encoding the CAR transgene and the other encoding the SB100x transposase) have antitumor activity *in vitro* and *in vivo*, while presenting long-term CAR expression. As a plasmid-based system, this method is less cumbersome and potentially has a reduced overall cost when compared to viral vectors. Moreover, different studies have shown that the transgene integration profile of SB vectors are close to random(19–21), which confers an advantage over retroviral and lentiviral vectors that have an integration profile biased towards transcriptionally-active sites(22–24).

Recently, Ghassemi et al reported that decreasing the time of *in vitro* T cell expansion increased its antitumor activity *in vivo*, and lower doses of CAR T cell were able to control leukemia growth (25). This report goes in line with the literature, showing that considerable T cell differentiation occur after *in vitro* expansion, with less differentiated, central memory-like T cells being associated with improved antitumor activity in preclinical models(26–28) and patients(29). In this proof-of-principle paper, we take this concept one step further and show that, by using SB transposon system and electroporation-based gene delivery, CAR-T cells can be generated and directly used for therapy, without the need of *ex vivo* activation and expansion protocols. We show that this point-of-care (POC) approach is efficient against two different B cell leukemia models (RS4;11 and Nalm-6), constituting a potential new method for the generation and application of CAR-T cell therapy.

## Material and Methods

### Plasmids

The 19BBz CAR sequence was supplied by Dr. Dario Campana (St Jude Children’s Research Hospital, Memphis, TN). The sequence was codon optimized and synthesized by Genscript (Piscataway, NJ) and a myc-tag was added between CD8α signal peptide and scFv to allow flow cytometry-based detection as described elsewhere (15). The transgene was cloned using AgeI/NotI sites in the transposon vector pT3 provided by Dr. Richard Morgan (NIH). The SB100x transposase which is encoded by plasmid pCMV-SB100x was provided by Dr Sang Won Han (Federal University of São Paulo [UNIFESP], Brazil).

### Cell lines and primary cells

Nalm-6 and RS4;11 were modified by lentivirus to express a fluorescent GFP protein as previously described(15) and were selected by fluorescence-activated cell sorting (FACS). These cells were cultivated RPMI (Gibco,CA) complete medium supplemented with 10% fetal calf serum, 2 mM L-Glu and 100 U/mL penicillin and 100 μg/mL streptomycin (Sigma-Aldrich, MO) at 37°C, 5% CO_2_. Peripheral blood mononuclear cells (PBMCs) were collected from blood bank of Brazilian National Cancer Institute (INCA). Healthy donors signed a board–approved informed consent. Mononuclear cells were isolated by density gradient centrifugation protocol (Ficoll-Hypaque-1077 (Sigma-Aldrich); slow acceleration/deceleration off; centrifugation for 20 min at 890g). Cells were washed three times with PBS. The use of PBMCs was approved by an institutional review board (Brazilian National Cancer Institute [INCA] Ethics Committee and performed in accordance with the Declaration of Helsinki.

### PBMCs electroporation and expansion

For PBMCs genetic modification, we used Lonza^®^ Nucleofector^®^ II electroporation system as described in Chicaybam, 2013 (30). Briefly, we transferred 10^7^ or 3×10^7^ cells with 100ul of 1SM buffer, 20ug of pT3-19BBz and 1ug SB100x plasmids into a 0.2 cm cuvette (Mirus Biotech^®^, Madison, WI) and cells were electroporated with the U-14 program. The mock condition was electroporated with SB100x plasmid only. Directly after electroporation, 1mL of RPMI medium supplemented with 2 mM L-glutamine, 20% fetal calf serum and IL-2 (50 U/mL; Proleukin, Zodiac) without antibiotic was added to the cuvette and cells were transferred to a 6-well plate. After 4 hours or 24 hours resting in RPMI with 20% fetal calf serum, cells were centrifuged, resuspended in PBS and used for POC condition. For the expansion experiments, cells were activated with T Cell TransAct CD3/CD28 (Miltenyi Biotec, Germany) at 1:200 concentration 24h after electroporation and the next day the supernatant was removed to withdraw beads. For the experiment referring to Figure 4, whole blood was incubated with RosetteSep™ Human T Cell Enrichment Cocktail (StemCell) for 15 min at the concentration of 30 μl to 1 ml of blood for purification of CD3+ cells. The cells were further processed as described above. For the *in vitro* experiment for Supplementary Fig. 1, T cells were cultured for 3 days with 200U IL-2 and cytotoxicity assays were performed as described below.

### Cytotoxicity Assay

This experiment was performed according to Neri et al,2001(31). Calcein-acetoxymethyl (Calcein-AM) (Thermo Fisher scientific,Burlington,CAN), was used to stain RS4;11 and Nalm-6 cells. The target cells were incubated in complete medium with 15 μM calcein-AM at final concentration of 10^6^/ml during 30min at 37°C and washed twice with medium. After this procedure, the target cells were resuspended at a final concentration of 5×10^3^ cells/50μl of medium. Effector cells were serially diluted from 50:1 to 0.7:1 of Effector:Target in a V-bottom plate and co-cultured with the 5×10^3^ target cells for 4 hours at 37°C and 5% CO2. After this procedure 75 ul of the supernatant were harvested and measured using a Tecan’s Infinite^®^ series of microplate readers (excitation filter: 485 ± 9 nm; band-pass filter: 530 ± 9 nm). The percent of lysis was calculated according to the formula: [(test release – spontaneous release)/(maximum release – spontaneous release)] × 100.

### Flow cytometry and antibodies

FACSCalibur (BD Biosciences) was used to perform the cell phenotype analysis. The cells previously washed with PBS were incubated with antibody for 15-20 min and then washed again. The following antibodies were used: anti-CD62L-FITC (Thermo Fisher Scientific Cat# 11-0629-42, RRID:AB_10667774), anti-CCR7-PE(Thermo Fisher Scientific Cat# 12-1979-42, RRID:AB_10670625), anti-CD3-PE-Cy5 (Thermo Fisher Scientific Cat# 15-0038-42, RRID:AB_10598354), anti-CD45RO-PE-Cy7 (Thermo Fisher Scientific Cat# 25-0457-42, RRID:AB_10718534), anti-CD14-FITC(Thermo Fisher Scientific Cat# 11-0149-42, RRID:AB_10597597), anti-CD56-PE (Thermo Fisher Scientific Cat# 12-0566-42, RRID:AB_2572563), anti-CD19-PercP Cy5.5 (Thermo Fisher Scientific Cat# 45-0199-42, RRID:AB_2043821), anti-CD19-FITC (Thermo Fisher Scientific Cat# 11-0199-42, RRID:AB_10669461), anti-Myc tag Alexa 647 and anti-Myc tag Alexa Fluor 488 (clone 9E10) (Santa Cruz Biotechnology Cat# sc-40, RRID:AB_627268), anti-CD4-PercP (BioLegend Cat# 344624, RRID:AB_2563326), anti-CD8-APC-Cy7 (BioLegend Cat# 344714, RRID:AB_2044006). Data was analyzed using the FlowJo software v10.1 (FlowJo, RRID:SCR_008520)

### Xenograft model

We used eight- to twelve–week-old NOD-SCID IL2R gamma null (NSG) mice. 5×10^6^ RS4;11 GFP or 10^5^ Nalm-6 GFP were inoculated intravenously on the tail vein. Following this procedure, the animals were randomized by weight to ensure no inoculation bias. The number of injected cells varied according to the experiment and is indicated throughout the results. PBS group indicates animals treated with saline (tumor-only control group) on the same day as treatment with 19BBz or mock cells. During the experiment, the blood was collected by facial vein using a 5 mm lancet (Goldenrod) and blood was placed in a tube with 0.5 M EDTA. Animal welfare was monitored, and animals euthanized when recommended in a CO_2_ chamber. The organs were then macerated, the red blood cells lysed with Ammonium-Chloride-Potassium (ACK) Lysing Buffer for 20 min and washed with PBS for flow cytometry analysis. The NSG mice were obtained from Jackson laboratories and all animal procedures were approved by the animal ethics committee of the Brazilian National Cancer Institute.

### Statistical analysis

The data was analyzed using Prism version 7.0 (GraphPad Prism, RRID:SCR_002798). Results of survival curve were analyzed using log-rank test and the p-value was described in each figure. For organs analysis were used Mann–Whitney test for pairwise comparisons. When using asterisks, consider *p < 0.05, **p < 0.01, ***p < 0.001.

## Results

### Evaluation of the potential antileukemic effect of the point-of-care approach

Point of care approaches have the potential to simplify and broaden CAR-T based therapies. In order to demonstrate the feasibility of this approach, we validated this strategy *in vivo* in preclinical models. First, we validated POC based protocol ability to restrain leukemia growth by injecting 5×10^6^ RS4;11 GFP cells in NSG mice on d+0, as demonstrated at the timeline (Fig. 1A). Three days later, PBMC from a healthy donor were isolated and electroporated with the pT3-19BBz plasmid (anti-CD19 CAR with 41BB and CD3ζ domains) and SB100x (the transposase that mediates transgene integration). Cells were rested for 4 hours and then 10^7^ total cells were inoculated to treat each mouse. After 24 hours of electroporation, we evaluated CAR expression by myc-tag detection *in vitro.* A total of 6,2×10^5^ CAR-T cells were injected into each animal (Fig. 1B).

**Figure 1:**
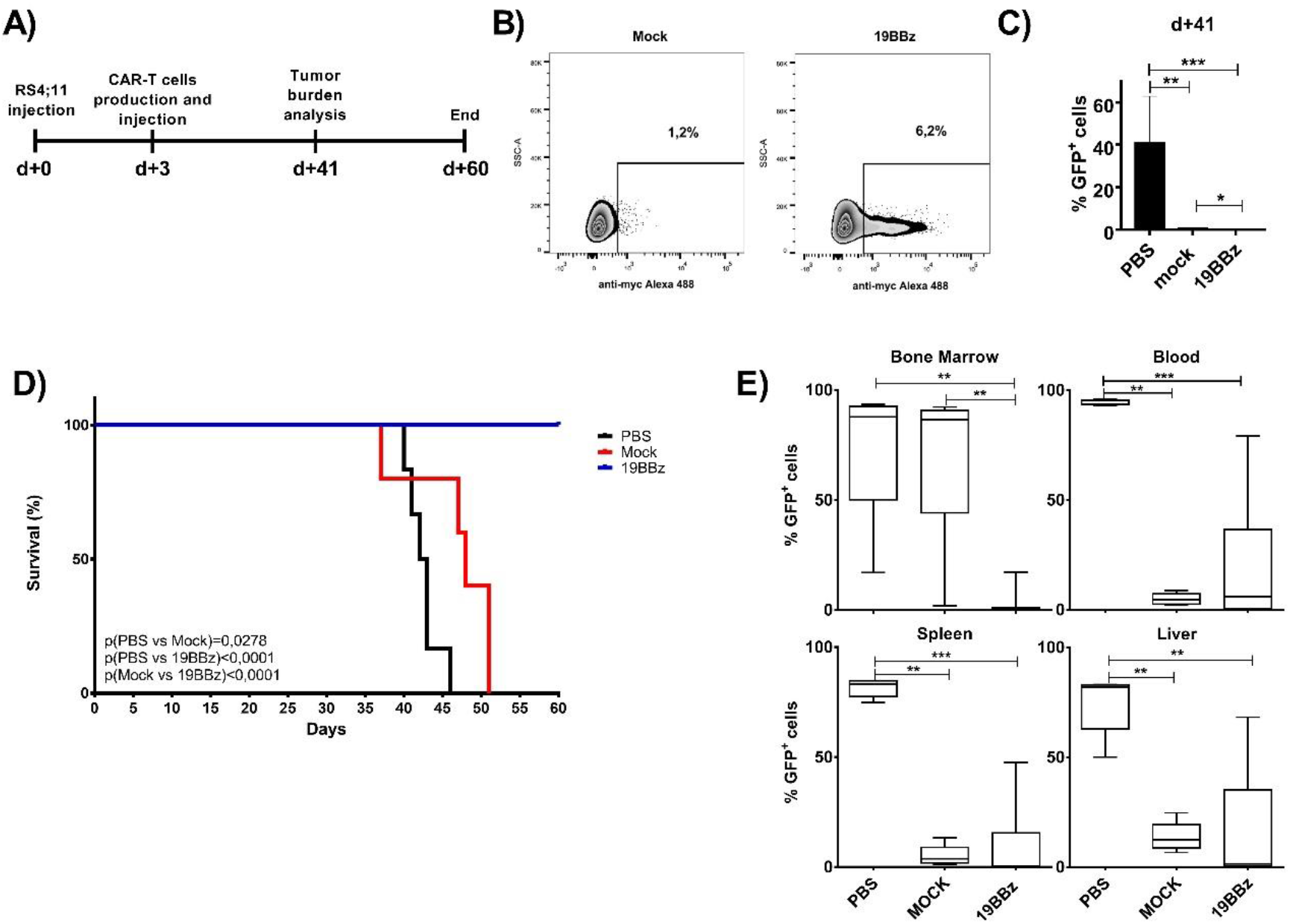
Evaluation of the potential antileukemic effect of the point-of-care approach. (A) Timeline of the experiment. NSG mice were inoculated with RS4;11 GFP tumor cells and, after 3 days, treated with CAR T cells electroporated 4 hours before treatment. (B) Expression of 19BBz CAR in T cells 24 hours after electroporation. Mock condition represents the electroporation of PBMC without 19BBz plasmid. (C) Animal blood was collected on day 41 for analysis of tumor burden of RS4;11 GFP by flow cytometry. (D) Kaplan-Meier plot of survival data (PBS n=6; Mock n=5; 19BBz n=9). (E) After euthanasia, tumor burden in blood, bone marrow, spleen and liver were analyzed by flow cytometry. The survival curve was analyzed by log-rank test and for organ analysis the Mann-Whitney test was used for paired comparisons. Consider * p <0.05, ** p <0.01, *** p <0.001.

During the experiment, we analyzed tumor burden by measuring RS4;11 GFP expression in the blood of animals over time (Fig. 1C). The group that received 19BBz CAR-T cells showed a decreased tumor burden in blood when compared to PBS treated mice (p=0.0004). Mock and 19BBz groups showed a lower statistical difference in blood leukemia burden (p=0.0196), suggesting an antitumor activity by untransfected cells. This lower tumor burden also impacted the survival curve (Fig. 1D), demonstrating an improvement in mock group when compared with PBS treated mice (p=0.0278).

Interestingly, 19BBz cells were able to greatly improve the overall survival of mice when compared to mock cells. At day 60, all 19BBz animals remained alive and were euthanized, the tumor burden of organs was then checked by flow cytometry and compared between groups (Fig. 1E). PBS group presented a high tumor burden in all organs analyzed and no significant difference was observed between mock and 19BBz, except for bone marrow, where 19BBz treated mice showed lower tumor burden.

The survival curve and bone marrow tumor burden indicate the effectiveness potential of the POC approach. Cells used 4 hours after electroporation do not yet express the CAR molecule, rendering impossible to evaluate the number of CAR-T cells infused in advance (such evaluation can only be performed at least 24h post gene transfer).

We usually evaluated CAR expression 24 hours after electroporation and, when we evaluated 5 donors for 3 days *in vitro* by keeping T cells in minimal culture conditions (without activation), CAR expression percentages remained overall stable (Supplementary Fig. 1A-B), so we can assume flow cytometry evaluation of CAR expression at 24h is a bona fide readout for CAR-T cell number evaluation. The *in vitro* antitumor capacity of CAR-T cells was low or absent and was similar to mock cells in the cytotoxicity assay against RS4;11 and Nalm-6 in these three days after electroporation (Supplementary Fig. 1C). In addition, CAR expression was restricted to CD3+ cells, with low or absent CAR expression in CD19+ or CD56+ cells (Supplementary Fig. 2) Since the 24h rest enables CAR-T expression evaluation before injection, we decided to check for CAR-T cell antitumor potential after this short period of incubation.

### Comparison of anti-tumor activity of cells 4h vs or 24h post electroporation

For a better characterization of the cells being used in the treatment, we compared overall survival of animals treated with equivalent numbers of CAR-T cells kept in culture for 4h or 24h after the electroporation.

RS4;11 leukemia cells were engrafted in NSG animals and 3 days later CAR-T from 4h or 24h post electroporation from the same donor were injected in equivalent numbers for both groups (Fig. 2A). PBMC composition using CD14, CD19 and CD56 markers was similar in both groups (Fig. 2B), with most of the cells being T lymphocytes (CD14-CD19-CD56-). T cell memory populations were determined among CD4+ and CD8+ cells (Fig. 2C) showing a predominance of central memory (CM) cells in CD4+ lymphocytes and also the presence of effector memory (EM) and naïve cells. For the CD8+ population, the predominant cell subset is EM, with smaller contributions for CM and naïve cells. No major difference in composition could be observed when comparing mock and 19BBz conditions.

**Figure 2:**
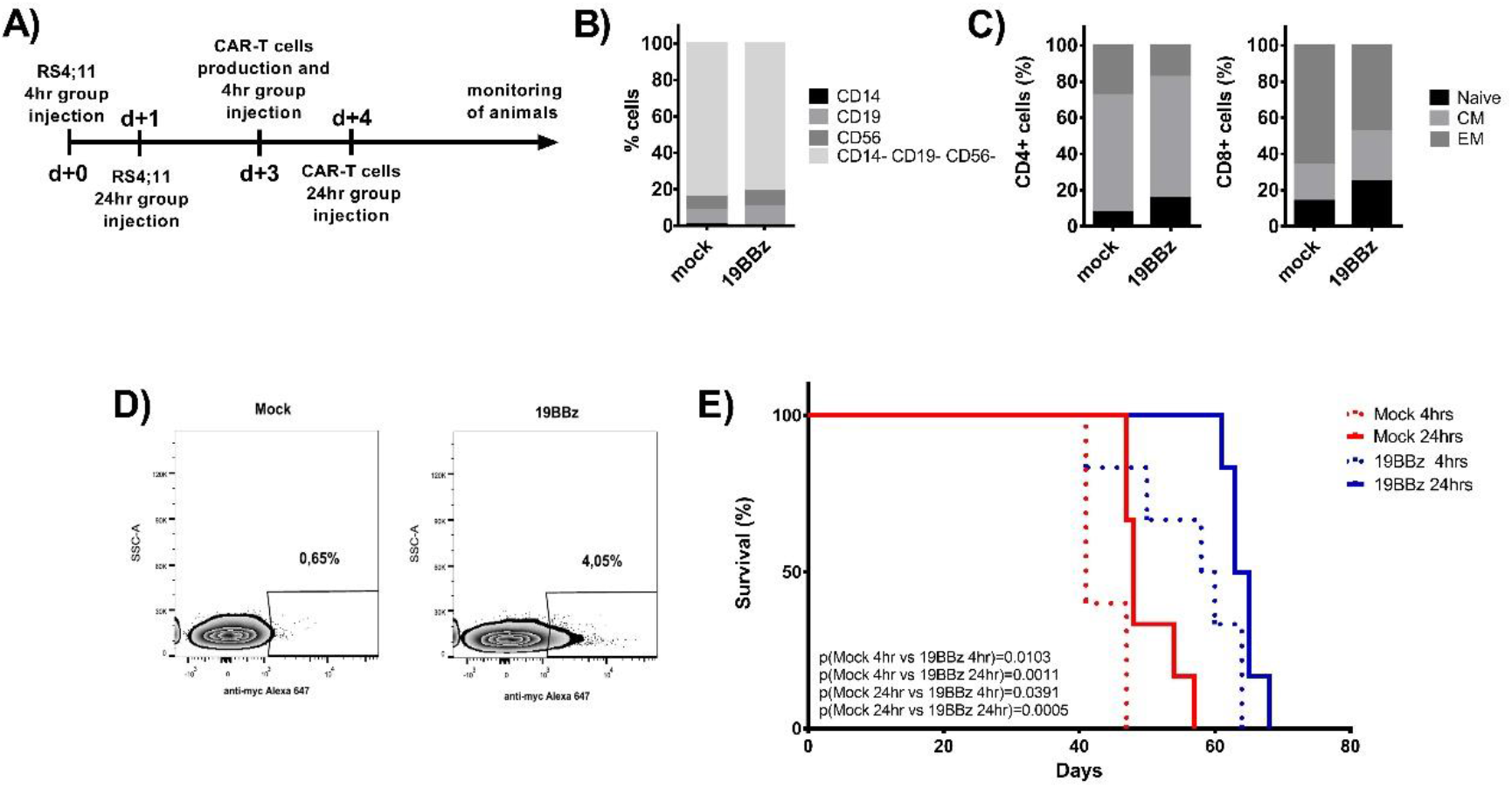
Comparison of antitumor activity in treated animals with cells electroporated ex4 or 24 hours earlier. (A) Timeline of the experiment. Animals were treated with CAR T cells from the same donor, but at different times after its production: 4 or 24 hours after electroporation. (B) Immunophenotypic characterization of cells and (C) memory phenotype characterization evaluated in CD4+ and CD8+ T cells and represented by Naive (CD45RO-), Central Memory (CM, CD45RO+ and presence of either CD62L or CCR7 markers) and effector Memory (CD45RO+CD62L-CCR7-) 24 hours after electroporation. (D) Expression of 19BBz CAR in T cells 24 hours after electroporation. (E) Kaplan-Meier plot of overall survival data (Mock 4hrs n=5; Mock 24hrs n=6; 19BBz 4hrs n=6; 19BBz 24hrs n=6). The survival curve was analyzed by log-rank test.

We adjusted the number of total cells injected to 10^6^ for all groups. After 24 hours of electroporation, the CAR expression could be detected in about 4% of the cells, meaning that 4×10^4^ CAR-T positive cells were injected (Fig. 2D). All the 19BBz conditions outperformed the corresponding mock conditions, both for 4h and 24h resting cells (Fig. 2E). When comparing 4h vs 24h between mock or 19BBz conditions, 24h resting seems to present a slight tendency to provide better survival, although statistical significance was never reached.

The RS4;11 model represents a valuable leukemia model that can reach very high tumor load in the periphery by day 21 and still allow the animals to survive for a few weeks, killing the mice by day 45 (Fig. 1D). In order to validate these results in a more aggressive mouse model, we tested the same approach in NSG mice engrafted with Nalm-6 leukemia, which leads to animal death by day 21.

### Effectiveness of the point-of-care approach in animals engrafted with Nalm-6

The Nalm-6 cell line is widely used and is a well-established model in several works with CAR-T cell therapies(32–34). We injected 10^5^ Nalm-6 per animal and treated the groups with CAR-T cells two days later (Fig. 3A). Using the 24h POC CAR-T generation approach, 18,5% of the cells showed CAR expression 24h after electroporation (Fig. 3B) and a total of 7×10^5^CAR-T cells were injected into each animal. The 19BBz treated group largely outperformed both PBS and mock treated groups in survival analysis (Fig. 3C). PBS and mock groups showed no difference in survival curve, indicating that mock electroporated cells have no antitumor activity in this model. After euthanasia, the 19BBz group showed lower tumor burden in all the evaluated organs when compared with the other groups, except in bone marrow. This inversion of tumor burden may be due to the longer survival time of the 19BBz group that allowed the accumulation of tumor cells in this organ. (Fig. 3D).

**Figure 3:**
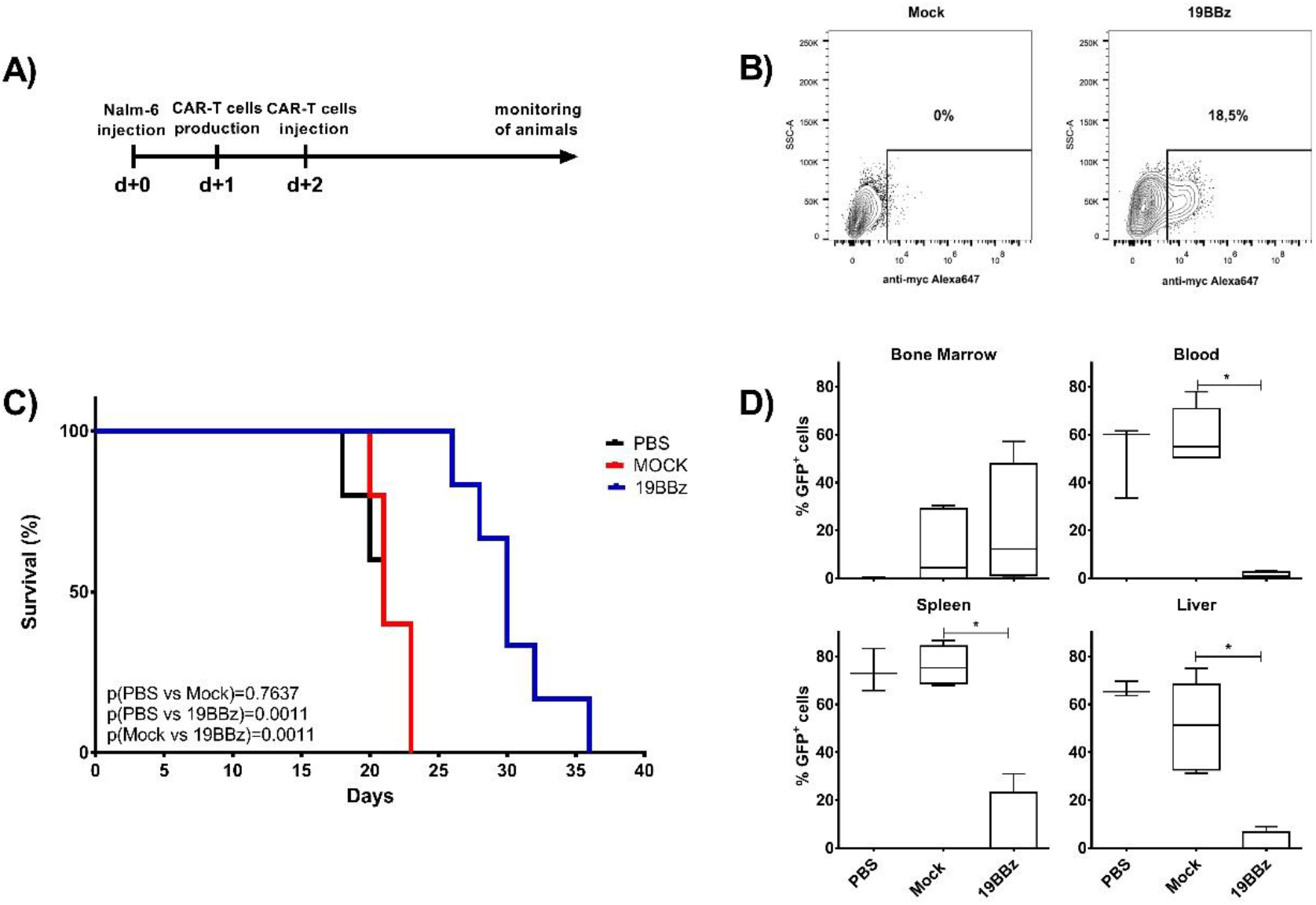
Effectiveness of point-of-care (POC) approach in animals engrafted with Nalm-6. (A) Timeline of the experiment. NSG mice were inoculated with 10^5^ Nalm-6 GFP cells and were treated after 2 days with CAR T cells produced 24 hours earlier. (B) Expression of 19BBz CAR in T cells 24 hours after electroporation. (C) Kaplan-Meier plot of survival data (PBS n=5; Mock n=5; 19BBz n=6). (D) After euthanasia, tumor burden of blood, bone marrow, spleen and liver were analyzed by flow cytometry. The survival curve was analyzed by log-rank test and for organ analysis the Mann-Whitney test was used for paired comparisons. Consider * p <0.05, ** p <0.01, *** p <0.001.

### Antitumor activity of Point-of-care and anti CD3/CD28 beads expanded cells

If a POC approach is to be used clinically, it should perform at least as well as the current bead-based expansion protocols. Hence, we conducted experiments to compare both approaches side-by-side. We divided the animals into two groups, POC and Expansion and used the same donor as source of T cells, adjusting the dose of CAR-T cells infused to be equal for both conditions.

On this occasion we purified T cells from PBMCs to ensure no antitumor effect by natural killer (NK) cells and ensure cell composition homogeneity in both groups by injecting only CD3+ cells. As previously described by our group, NK cells may have accounted for improved survival of the mock group compared to PBS treated animals when the infused product contains this lymphocyte population (15). T cells were electroporated and infused into the animals after 24 hours in the POC group. A fraction of the cells from the same electroporation were stimulated with CD3/CD28 Transact beads and expanded for 8 days. Figure 4A shows the expansion of these cells in culture, with 19BBz cells displaying higher expansion capabilities when compared to the mock electroporated cells. As expected, the memory cell profiles changed over the *in vitro* culture period. Only low levels of CD4+ and CD8+ Naive T cells could be detected after 8 days of culture and the lymphocytes consisted basically of CM cells at the end of the culture for both CD4 and CD8 subsets (Fig. 4B).

**Figure 4:**
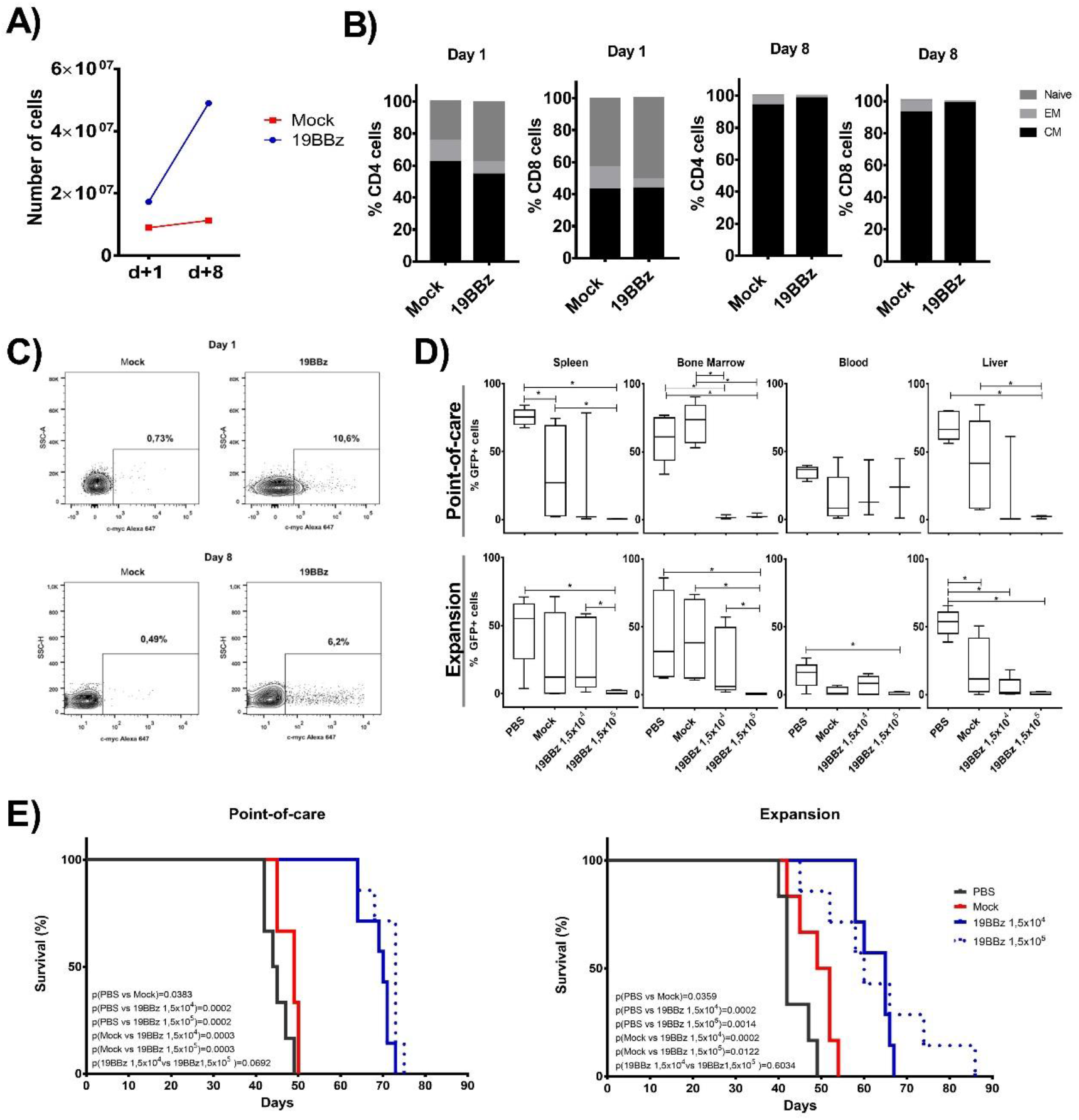
Comparison of the efficiency of antitumor activity between the Point-of-care approach and cells expanded with anti-CD3/CD28 coated beads. (A) Comparison of the expansion of CAR+ cells. Cells from this donor were used to treat NSG mice engrafted with RS4;11 in the POC or expanded cells approach. (B) Memory phenotype was evaluated at day 1 and day 8 after electroporation in CD4+ and CD8+ T cells and were represented by Naive (CD45RO-), Central Memory (CM, CD45RO+ and presence of either CD62L or CCR7 markers) and Effector Memory (CD45RO+CD62L-CCR7-). (C) Expression of 19BBz CAR in T cells 24 hours after electroporation and at day 8 of the expansion protocol. (D) Evaluation by flow cytometry of the tumor burden of the organs in the groups after euthanasia. (E) Kaplan-Meier plot of survival data (POC group: PBS n=6; Mock n=6; 19BBz 1,5×10^4^ n=7; 19BBz 1,5×10^5^ n=7. Expansion group: PBS n=6; Mock n=6; 19BBz 1,5×10^4^ n=7; 19BBz 1,5×10^5^ n=7). The survival curve was analyzed by log-rank test and for organ analysis the Mann-Whitney test was used for paired comparisons. Consider * p <0.05, ** p <0.01, *** p <0.001.

CAR expression was evaluated by cytometry to match the same number of CAR-T cells in POC and Expansion treated groups (Fig. 4C). We tested two different CAR-T cell numbers, injecting 1,5×10^5^ or 1,5×10^4^ CAR+ cells in two separate groups. Mock condition animals received 1,4×10^6^ total cells. We analyzed the percentage of GFP cells in spleen, bone marrow, blood and liver after euthanasia and the group that received the largest number of CAR-T cells showed the lowest tumor burden in almost all cases in both the POC and Expansion groups (Fig. 4D).

Survival of animals from the POC group proved to be similar to that of the Expansion group (Fig. 4E). In both cases, all animals receiving any dose of CAR-T cells had improved survival. No consistent dose-response effect was observed between 19BBz groups: POC p(1,5×10^4^ vs 1,5×10^5^ 19BBz) = 0,0692 and Expansion p(1,5×10^4^ vs 1,5×10^5^ 19BBz) = 0,6034.

This experiment was repeated using similar conditions. In this case, cells were expanded for 12 days and no change in CAR-T cell percentage was observed (Supplementary Fig. 3A). Central memory cells were also the major population after expansion (Supplementary Fig. 3B). Survival of animals from the POC group proved to be similar to that of the Expansion group using the same CAR-T cell dose. In this model, a higher dose of CAR-T cell confers a longer overall survival from both POC and Expansion groups (Supplementary Fig. 3C). Our data indicates that the antitumor capacity of the POC approach is comparable to the potency of the bead-based expansion protocol, indicating that POC based CAR therapy can be proposed as a clinical strategy.

## Discussion

CAR-T cells are now being successfully used for the treatment of B cell leukemias and lymphomas, constituting a new treatment option for these malignancies. Further developments in target antigen selection, CAR design and combination therapy are likely to extend CAR-T cell therapy to solid tumors(35,36). Thus, large scale production capabilities need to be developed in order to meet this expected increase in demand in the next years.

Current methods used for autologous CAR-T cell manufacturing rely on viral vector transduction and *ex vivo* expansion protocols to generate the therapeutic product, procedures that are difficult to scale up and that require skilled technicians. In an attempt to optimize production, companies opted for centralized production facilities that supply CAR-T cell therapies to qualified treatment centers, but this also imposes a considerable logistic challenge that contributes to the complexity of this treatment modality(37). Indeed, 7 of 88 patients enrolled in the ELIANA trial could not be treated with tisagenlecleucel due to manufacturing failures(38). Moreover, some patients with advanced disease may succumb to the disease prior to receiving the therapy. These aspects, combined with long (12-20 days) production time and high costs, constitute important bottlenecks for the widespread use of this technology. The use of automated CAR-T cell production equipments (for example, Miltenyi Prodigy) will likely improve the process, but the high upfront costs of acquisition and lack of customization might hamper their use in the short term(39,40).

Meanwhile, preclinical and clinical data is showing that T cells with a less differentiated phenotype, such as naïve or central memory T cells, have improved antitumor responses *in vivo*(41–43). In fact, a recent report described that the presence of a memory-like CD27+CD45RO-CD8+ T cell subpopulation was associated with improved responses in patients with chronic lymphocytic leukemia treated with anti-CD19 CAR-T cells (27). However, current T cell manufacturing protocols are not optimized to generate or expand these cell types and generally induce T cell differentiation, mainly related to exposure to cytokines like IL-2(44,45). Importantly, it was recently shown that producing CAR-T cells using a shorter (3-5 days) expansion protocol can induce less T cell differentiation, reducing the expression of inhibitory receptors and improving effector function(41,42). Importantly, it was recently shown that producing CAR-T cells using a shorter (3-5 days) expansion protocol can induce less T cell differentiation, reducing the expression of inhibitory receptors and improving effector function(25).

In this context, the use of SB system associated to nucleofection provides a compelling approach for the fast generation of minimally manipulated CAR-T cells. Being a plasmid-based, integrative genetic modification tool, this system can be delivered by electroporation and generate large numbers of CAR-T cells in a short time, consistent with a POC approach. As we showed in this report, T cells electroporated with 19BBz CAR and injected after only 4h induce increased survival of mice engrafted with RS4;11 B cell leukemia when compared to mock T cells (Fig. 1). A similar result was obtained when 19BBz were rested in culture for 24h after electroporation, demonstrating that 4×10^4^ CAR+ T cells are already sufficient to induce an increased survival in mice engrafted with RS4;11 B cell leukemia (Fig. 2). This is important since, for regulatory reasons, the 24h cultivation would allow rapid tests for sterility to be performed and minimal phenotypic characterization of the cells, such as CAR detection on T lymphocytes and dose adjustments to reduce potential toxicities. Supporting this idea, our data show that CAR expression at d+1 post nucleofection represents roughly the peak of CAR expression (Supplementary Fig. 1) and can be used as a proxy for CAR-T cell infusion doses. Our results also indicate that CAR-T cells at d+1 post CAR gene transfer do not display significant leukemia lysis capabilities *in vitro*, reinforcing the notion that these minimally manipulated CAR-T cells will expand and differentiate *in vivo*, acquiring their final phenotype and cytotoxic functions.

Our experiments also demonstrated that 19BBz CAR-T used in POC approach showed antitumor responses in a more aggressive B cell leukemia model using the Nalm-6 cell line. As shown in figure 3, 19BBz CAR-T cells induced improved overall survival and decreased tumor burden in blood, spleen and liver when compared to mock T cells. It is important to note that 7×10^5^ CAR-T cell were required to induce this response, as numbers lower than this were not effective *in vivo* (data not shown). Finally, side by side comparison of CAR-T cells generated after 8 days of *ex vivo* expansion with CAR-T cells used with POC approach showed equivalent efficiency *in vivo* in terms of percentage of tumor cells in organs (Fig. 4D) and overall survival (Fig. 4E), despite showing different distribution of T cell memory subpopulations in the cell preparations (Fig. 4B and Supplementary Fig. 3).

The results presented in this paper demonstrate the feasibility of the POC approach. The use of large-scale, closed electroporation systems, like Lonza Nucleofector 4D LV unit or Maxcyte’s electroporator, can process large quantities (up to 20×10^9^ cells per run) of PBMCs(46) and generate minimally-manipulated CAR-T cells in a short time, being compatible with GMP procedures. It was recently reported that 1.18 – 298×10^9^ PBMCs (median: 8.83×10^9^ cells) can be recovered after apheresis of pediatric and young adult patients with B-ALL or neuroblastoma enrolled in CAR clinical trials(47). A second paper reported recoveries ranging 0.12-100×10^9^ of PBMCs (median: 10.5×10^9^) in adult patients(48). Assuming 50% viability and 10% transfection efficiency, by processing around 3×10^9^ PBMCs it would be possible to generate 15×10^7^ CAR+ T cells, equivalent to 2.1×10^6^ CAR+ T cells/kg for a 70kg patient. Therefore, production of CAR-T cells for POC approach using currently available devices is potentially feasible and compatible with T cell doses used in the clinical practice. Nonetheless, the minimal manipulation of our POC approach has the potential to generate cells in better shape to provide leukemia protection *in vivo*, so the minimal doses required for *in vivo* function in patients are still to be determined.

The proposed approach in this report can be further potentiated with additional manipulations of T cells such as the addition of cytokines or immune checkpoint blockade secretion by T cells, as already described(49). In any case, the absence of an *ex vivo* expansion phase simplifies the CAR-T manufacturing process and has the potential to drastically decrease the overall cost, facilitating the adoption of this new therapy, especially in developing countries.

## References

1. Park JH, Rivière I, Gonen M, Wang X, Sénéchal B, Curran KJ, et al. Long-Term Follow-up of CD19 CAR Therapy in Acute Lymphoblastic Leukemia. N Engl J Med. 2018;378:449–59.

2. Maude SL, Laetsch TW, Buechner J, Rives S, Boyer M, Bittencourt H, et al. Tisagenlecleucel in Children and Young Adults with B-Cell Lymphoblastic Leukemia. N Engl J Med. 2018;378:439–48.

3. Neelapu SS, Locke FL, Bartlett NL, Lekakis LJ, Miklos DB, Jacobson CA, et al. Axicabtagene Ciloleucel CAR T-Cell Therapy in Refractory Large B-Cell Lymphoma. N Engl J Med. 2017;377:2531–44.

4. Schuster SJ, Bishop MR, Tam CS, Waller EK, Borchmann P, McGuirk JP, et al. Tisagenlecleucel in Adult Relapsed or Refractory Diffuse Large B-Cell Lymphoma. N Engl J Med. 2019;380:45–56.

5. June CH, Sadelain M. Chimeric Antigen Receptor Therapy. N Engl J Med. 2018;379:64–73.

6. Fry TJ, Shah NN, Orentas RJ, Stetler-Stevenson M, Yuan CM, Ramakrishna S, et al. CD22-targeted CAR T cells induce remission in B-ALL that is naive or resistant to CD19-targeted CAR immunotherapy. Nat Med. 2018;24:20–8.

7. Pan J, Niu Q, Deng B, Liu S, Wu T, Gao Z, et al. CD22 CAR T-cell therapy in refractory or relapsed B acute lymphoblastic leukemia. Leukemia. 2019;1–13.

8. Xu J, Chen L-J, Yang S-S, Sun Y, Wu W, Liu Y-F, et al. Exploratory trial of a biepitopic CAR T-targeting B cell maturation antigen in relapsed/refractory multiple myeloma. Proc Natl Acad Sci. 2019;116:9543–51.

9. Cohen AD, Garfall AL, Stadtmauer EA, Melenhorst JJ, Lacey SF, Lancaster E, et al. B cell maturation antigen–specific CAR T cells are clinically active in multiple myeloma. J Clin Invest. 2019;129:2210–21.

10. Brudno JN, Maric I, Hartman SD, Rose JJ, Wang M, Lam N, et al. T Cells Genetically Modified to Express an Anti–B-Cell Maturation Antigen Chimeric Antigen Receptor Cause Remissions of Poor-Prognosis Relapsed Multiple Myeloma. J Clin Oncol. 2018;36:2267–80.

11. O’Rourke DM, Nasrallah MP, Desai A, Melenhorst JJ, Mansfield K, Morrissette JJD, et al. A single dose of peripherally infused EGFRvIII-directed CAR T cells mediates antigen loss and induces adaptive resistance in patients with recurrent glioblastoma. Sci Transl Med. 2017;9:eaaa0984.

12. Wang X, Rivière I. Clinical manufacturing of CAR T cells: foundation of a promising therapy. Mol Ther Oncolytics. 2016;3:16015.

13. Poorebrahim M, Sadeghi S, Fakhr E, Abazari MF, Poortahmasebi V, Kheirollahi A, et al. Production of CAR T-cells by GMP-grade lentiviral vectors: Latest advances and future prospects. Crit Rev Clin Lab Sci. 2019;0:1–27.

14. Vormittag P, Gunn R, Ghorashian S, Veraitch FS. A guide to manufacturing CAR T cell therapies. Curr Opin Biotechnol. 2018;53:164–81.

15. Chicaybam L, Abdo L, Carneiro M, Peixoto B, Viegas M, de Sousa P, et al. CAR T Cells Generated Using Sleeping Beauty Transposon Vectors and Expanded with an EBV-Transformed Lymphoblastoid Cell Line Display Antitumor Activity In Vitro and In Vivo. Hum Gene Ther. 2019;30:511–22.

16. Magnani CF, Mezzanotte C, Cappuzzello C, Bardini M, Tettamanti S, Fazio G, et al. Preclinical Efficacy and Safety of CD19CAR Cytokine-Induced Killer Cells Transfected with Sleeping Beauty Transposon for the Treatment of Acute Lymphoblastic Leukemia. Hum Gene Ther. 2018;29:602–13.

17. Monjezi R, Miskey C, Gogishvili T, Schleef M, Schmeer M, Einsele H, et al. Enhanced CAR T-cell engineering using non-viral *Sleeping Beauty* transposition from minicircle vectors. Leukemia. 2017;31:186–94.

18. Kebriaei P, Izsvák Z, Narayanavari SA, Singh H, Ivics Z. Gene Therapy with the Sleeping Beauty Transposon System. Trends Genet. 2017;33:852–70.

19. Huang X, Guo H, Tammana S, Jung Y-C, Mellgren E, Bassi P, et al. Gene Transfer Efficiency and Genome-Wide Integration Profiling of Sleeping Beauty, Tol2, and PiggyBac Transposons in Human Primary T Cells. Mol Ther. 2010;18:1803–13.

20. Yant SR, Wu X, Huang Y, Garrison B, Burgess SM, Kay MA. High-Resolution Genome-Wide Mapping of Transposon Integration in Mammals. Mol Cell Biol. 2005;25:2085–94.

21. Jong J de, Akhtar W, Badhai J, Rust AG, Rad R, Hilkens J, et al. Chromatin Landscapes of Retroviral and Transposon Integration Profiles. PLOS Genet. 2014;10:e1004250.

22. Field A-C, Vink C, Gabriel R, Al-Subki R, Schmidt M, Goulden N, et al. Comparison of Lentiviral and Sleeping Beauty Mediated αβ T Cell Receptor Gene Transfer. PLOS ONE. 2013;8:e68201.

23. Wu X, Li Y, Crise B, Burgess SM. Transcription Start Regions in the Human Genome Are Favored Targets for MLV Integration. Science. 2003;300:1749–51.

24. Qin D-Y, Huang Y, Li D, Wang Y-S, Wang W, Wei Y-Q. Paralleled comparison of vectors for the generation of CAR-T cells. Anticancer Drugs. 2016;27:711–22.

25. Ghassemi S, Nunez-Cruz S, O’Connor RS, Fraietta JA, Patel PR, Scholler J, et al. Reducing Ex Vivo Culture Improves the Antileukemic Activity of Chimeric Antigen Receptor (CAR) T Cells. Cancer Immunol Res. 2018;6:1100–9.

26. Gattinoni L, Klebanoff CA, Palmer DC, Wrzesinski C, Kerstann K, Yu Z, et al. Acquisition of full effector function in vitro paradoxically impairs the in vivo antitumor efficacy of adoptively transferred CD8^+^ T cells. J Clin Invest. 2005;115:1616–26.

27. Klebanoff CA, Gattinoni L, Torabi-Parizi P, Kerstann K, Cardones AR, Finkelstein SE, et al. Central memory self/tumor-reactive CD8+ T cells confer superior antitumor immunity compared with effector memory T cells. Proc Natl Acad Sci. 2005;102:9571–6.

28. Alizadeh D, Wong RA, Yang X, Wang D, Pecoraro JR, Kuo C-F, et al. IL15 Enhances CAR-T Cell Antitumor Activity by Reducing mTORC1 Activity and Preserving Their Stem Cell Memory Phenotype. Cancer Immunol Res. 2019;7:759–72.

29. Fraietta JA, Lacey SF, Orlando EJ, Pruteanu-Malinici I, Gohil M, Lundh S, et al. Determinants of response and resistance to CD19 chimeric antigen receptor (CAR) T cell therapy of chronic lymphocytic leukemia. Nat Med. 2018;24:563–71.

30. Chicaybam L, Sodre AL, Curzio BA, Bonamino MH. An efficient low cost method for gene transfer to T lymphocytes. PloS One. 2013;8:e60298.

31. Neri S, Mariani E, Meneghetti A, Cattini L, Facchini A. Calcein-acetyoxymethyl cytotoxicity assay: standardization of a method allowing additional analyses on recovered effector cells and supernatants. Clin Diagn Lab Immunol. 2001;8:1131–5.

32. Brentjens RJ, Santos E, Nikhamin Y, Yeh R, Matsushita M, La Perle K, et al. Genetically targeted T cells eradicate systemic acute lymphoblastic leukemia xenografts. Clin Cancer Res Off J Am Assoc Cancer Res. 2007;13:5426–35.

33. Castella M, Boronat A, Martín-Ibáñez R, Rodríguez V, Suñé G, Caballero M, et al. Development of a Novel Anti-CD19 Chimeric Antigen Receptor: A Paradigm for an Affordable CAR T Cell Production at Academic Institutions. Mol Ther Methods Clin Dev. 2018;12:134–44.

34. Feucht J, Sun J, Eyquem J, Ho Y-J, Zhao Z, Leibold J, et al. Calibration of CAR activation potential directs alternative T cell fates and therapeutic potency. Nat Med. 2019;25:82–8.

35. Chicaybam L, Sodré AL, Bonamino M. Chimeric antigen receptors in cancer immuno-gene therapy: current status and future directions. Int Rev Immunol. 2011;30:294–311.

36. Chicaybam L, Bonamino MH. Moving receptor redirected adoptive cell therapy toward fine tuning of antitumor responses. Int Rev Immunol. 2014;33:402–16.

37. Highfill SL, Stroncek DF. Overcoming Challenges in Process Development of Cellular Therapies. Curr Hematol Malig Rep. 2019;14:269–77.

38. US Food and Drug Administration. FDA briefing document: Oncologic Drugs Advisory Committee meeting; BLA 125646; Tisagenlecleucel, Novartis Pharmaceuticals Corporation. FDA https://www.fda.gov/downloads/AdvisoryCommittees/CommitteesMeetingMaterials/Drugs/OncologicDrugsAdvisoryCommittee/UCM566166.pdf (2017).

39. Zhu F, Shah N, Xu H, Schneider D, Orentas R, Dropulic B, et al. Closed-system manufacturing of CD19 and dual-targeted CD20/19 chimeric antigen receptor T cells using the CliniMACS Prodigy device at an academic medical center. Cytotherapy. 2018;20:394–406.

40. Zhang W, Jordan KR, Schulte B, Purev E. Characterization of clinical grade CD19 chimeric antigen receptor T cells produced using automated CliniMACS Prodigy system. Drug Des Devel Ther. 2018;12:3343–56.

41. Gattinoni L, Zhong X-S, Palmer DC, Ji Y, Hinrichs CS, Yu Z, et al. Wnt signaling arrests effector T cell differentiation and generates CD8^+^ memory stem cells. Nat Med. 2009;15:808–13.

42. Klebanoff CA, Scott CD, Leonardi AJ, Yamamoto TN, Cruz AC, Ouyang C, et al. Memory T cell–driven differentiation of naive cells impairs adoptive immunotherapy. J Clin Invest. 2016;126:318–34.

43. Busch DH, Fräβle SP, Sommermeyer D, Buchholz VR, Riddell SR. Role of memory T cell subsets for adoptive immunotherapy. Semin Immunol. 2016;28:28–34.

44. Kaartinen T, Luostarinen A, Maliniemi P, Keto J, Arvas M, Belt H, et al. Low interleukin-2 concentration favors generation of early memory T cells over effector phenotypes during chimeric antigen receptor T-cell expansion. Cytotherapy. 2017;19:689–702.

45. Raeber ME, Zurbuchen Y, Impellizzieri D, Boyman O. The role of cytokines in T-cell memory in health and disease. Immunol Rev. 2018;283:176–93.

46. Hung C-F, Xu X, Li L, Ma Y, Jin Q, Viley A, et al. Development of Anti-Human Mesothelin-Targeted Chimeric Antigen Receptor Messenger RNA-Transfected Peripheral Blood Lymphocytes for Ovarian Cancer Therapy. Hum Gene Ther. 2018;29:614–25.

47. Ceppi F, Rivers J, Annesley C, Pinto N, Park JR, Lindgren C, et al. Lymphocyte apheresis for chimeric antigen receptor T-cell manufacturing in children and young adults with leukemia and neuroblastoma. Transfusion (Paris). 2018;58:1414–20.

48. Tuazon SA, Li A, Gooley T, Eunson TW, Maloney DG, Turtle CJ, et al. Factors affecting lymphocyte collection efficiency for the manufacture of chimeric antigen receptor T cells in adults with B-cell malignancies. Transfusion (Paris). 2019;59:1773–80.

49. Chan T, Gallagher J, Cheng N-L, Carvajal-Borda F, Plummer J, Govekung A, et al. CD19-Specific Chimeric Antigen Receptor-Modified T Cells with Safety Switch Produced Under “Point-of-Care” Using the Sleeping Beauty System for the Very Rapid Manufacture and Treatment of B-Cell Malignancies. Blood. 2017;130:1324–1324.

