## Supplementary data for "Development of CAR-T Cell Therapy for B-ALL Using a Point-of-Care Approach"

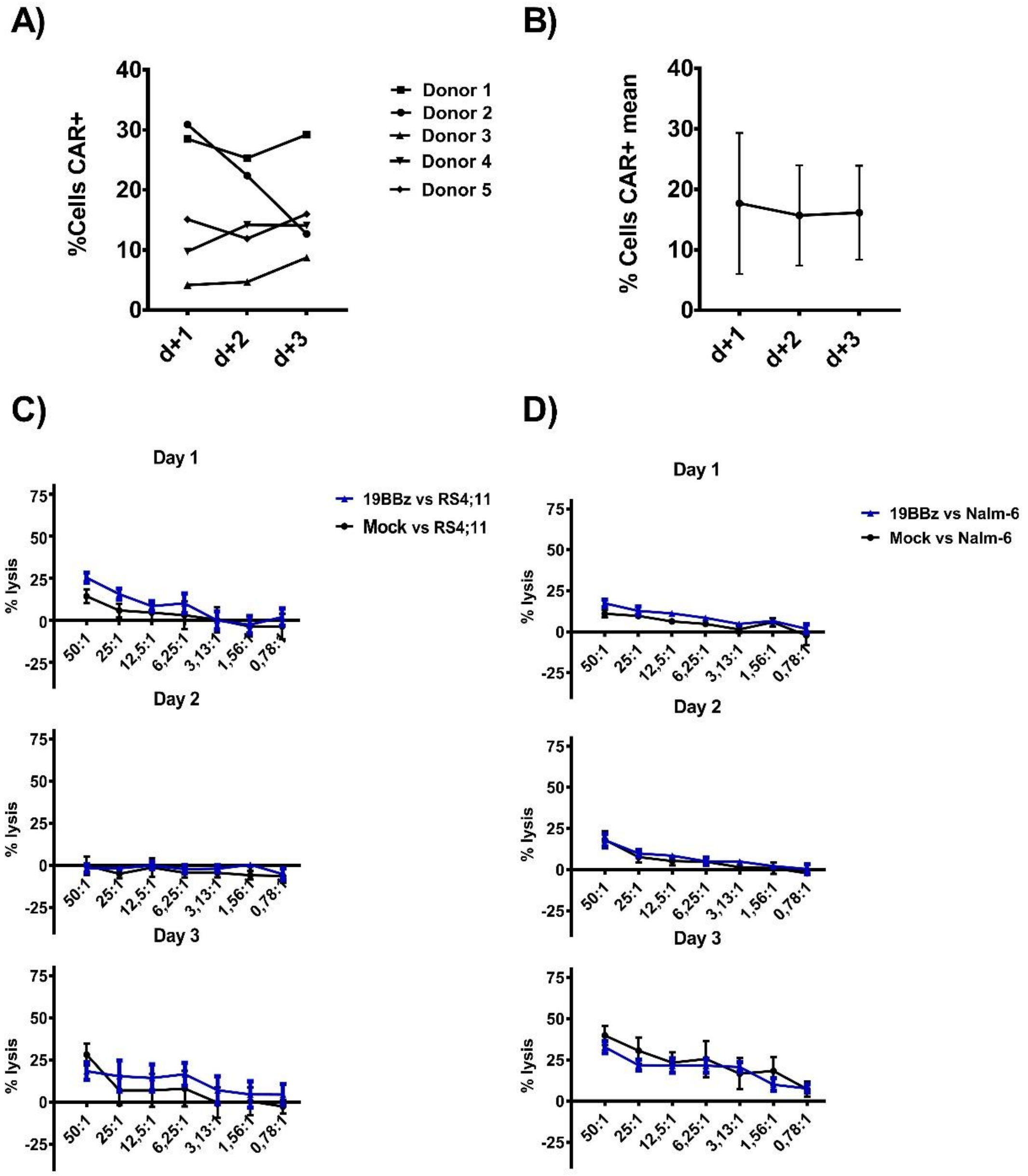


**Supplementary Figure 1:** Frequency of T cells expressing 19BBz CAR and *in vitro* cytotoxic activity evaluation. (A) Percentage of CAR-T cells at days 1, 2 and 3 after electroporation from five independent donors. T cells were maintained *in vitro* complete RPMI + 200U/mL IL-2 cultivation (without activation). (B) Mean CAR expression from donors depicted in (A). (C-D) Cytotoxicity assays were performed with the same 5 donors against RS4;11 and Nalm-6 in technical triplicate. Graphs represent the mean cytotoxic activity against RS4;11 (C) or Nalm-6 (D) from 5 donors at different Effector:Target rations at days 1, 2, and 3 after electroporation. Data is represented with mean +/-SEM.

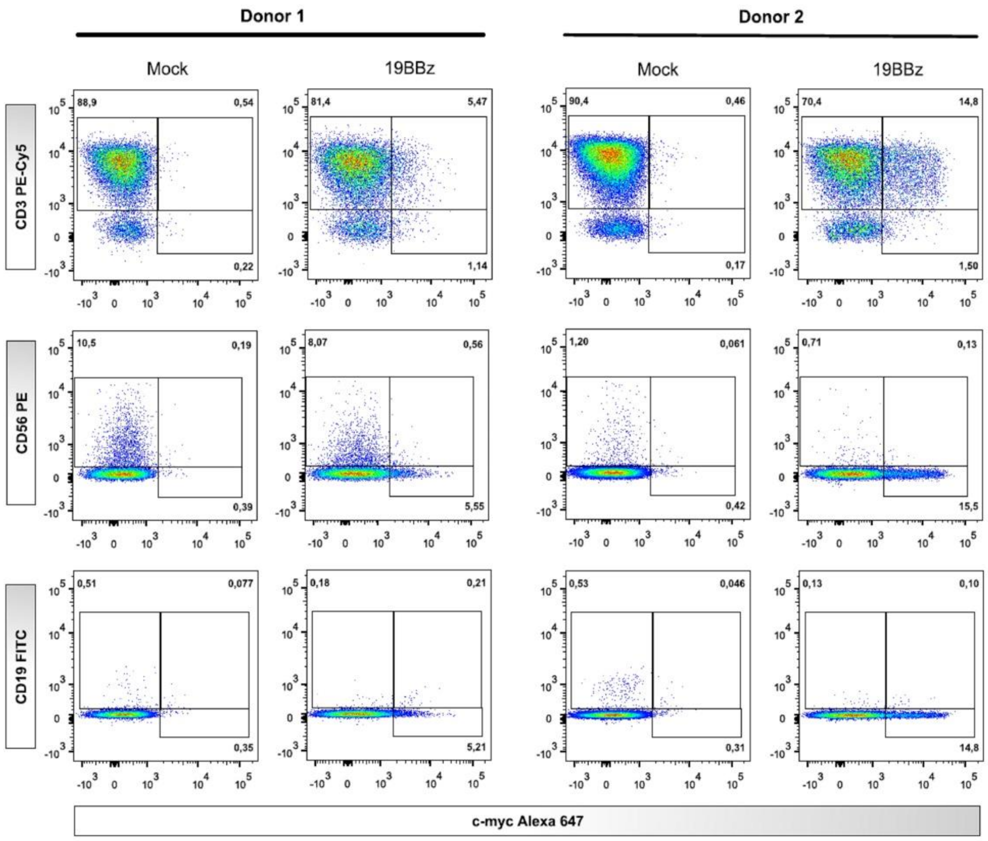


**Supplementary Figure 2:** CAR expression checked on CD3+, CD19+ and CD56+ cells. 24 hours after electroporation, the CAR expression was evaluated by flow cytometry by evaluating CD3, CD19 and CD56 markers.

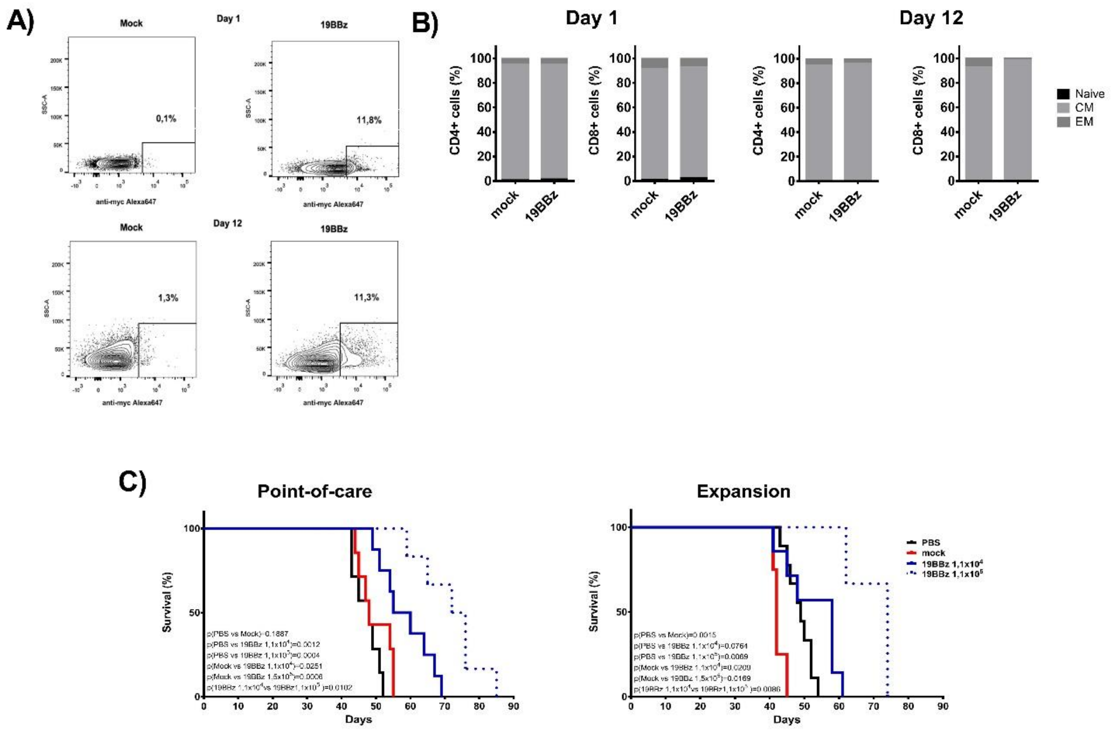


**Supplementary Figure 3:** Comparison of the efficiency of antitumor activity between the Point-of-care approach and expanded cells with anti-CD3/CD28 coated beads in RS4;11 tumor model. (A) Expression of 19BBz CAR in T cells 24 hours after electroporation and at day 12 of the beads-based expansion protocol. (B) Memory phenotype was evaluated at day 1 and day 12 in CD4+ and CD8+ T cells and represented by Naive (CD45RO-), Central Memory (CM, CD45RO+ and presence of either CD62L or CCR7 markers) and Effector Memory (CD45RO+CD62L-CCR7-). (C) Kaplan-Meier plot of survival data (POC group: PBS n=7; Mock n=7; 19BBz $1,1x10^4$ n=8; 19BBz $1,1x10^5$ n=6. Expansion group: PBS n=9; Mock n=4; 19BBz $1,1x10^4$ n=7; 19BBz $1,1x10^5$ n=3). The same donor was used to treat animals engrafted with RS4;11 in the POC approach and expanded cells strategies. The survival curve was analyzed by log-rank test.
